# Novel biochemical, structural and systems insights into inflammatory signaling revealed by contextual interaction proteomics

**DOI:** 10.1101/2021.09.18.460902

**Authors:** Rodolfo Ciuffa, Federico Uliana, Martin Mehnert, Cathy Marulli, Ari Satanowski, Pilar Natalia Rodilla Ramírez, Pascal Meier, Alessandro Ori, Matthias Gstaiger, Ruedi Aebersold

## Abstract

Protein-protein interactions (PPI) represent the main mode of the proteome organization in the cell. In the last decade, several large-scale representations of PPI networks have captured generic aspects of the functional organization of network components, but mostly lack the context of cellular states. However, the generation of contextual representations of PPI networks is essential for structural and systems-level modeling of biological processes and remains an unsolved challenge. In this study we describe an integrated experimental/computational strategy to achieve a contextualized modeling of PPI. This strategy defines the composition, stoichiometry, spatio-temporal organization and cellular requirements for the formation of target assemblies. We used this approach to generate an integrated model of the formation principles and architecture of a large signalosome, the TNF-receptor signaling complex (TNF-RSC). Overall, we show that the integration of systems- and structure-level information provides a generic, largely unexplored link between the modular proteome and cellular function.

**Significance Statement:** **In this work, we propose a critical shift in the way we analyze, and think the study of, protein-protein interactions (PPI)**, and present an experimental and computational framework to model them in the cellular context. We applied this framework to the signalosome **tumor necrosis factor receptor signaling complex (TNF-RSC)**, and generated an integrated model of its formation and architecture that provides new insights and resolved controversies regarding its organization and regulation. To achieve a **contextual modelling** of PPIs, we first optimized and developed, and then combined, approaches to map the **composition** of a target complex, its **absolute stoichiometry**, its **spatial organization** and assembly/disassembly **dynamics**, its temporal dependence on signaling, and its reliance on **cellular resources**.

## Introduction

As a consequence of the emergence of disruptive technologies, most classes of biomolecules can now be systematically measured. Typically, such OMIC technologies are initially used to generate generic, decontextualized maps of the respective molecular class, followed by subsequent studies that aim at relating the respective “OME” to the context of specific cell types or cellular states. This trajectory, exemplified by advances in genomics, transcriptomics, or proteomics has proven invaluable for basic and translational research.

The application of OMIC technologies to the study of PPIs has shown that protein assemblies mediate most biological functions (*1*). Such assemblies, whether stable or transient, can undergo dramatic changes in their composition, topology and activity as a function of the cellular state. To date, substantial progress has been made to generate generic maps of protein complexes and protein-protein interactions (*2,3,4*). These maps are essential to drive discoveries and define general properties of the proteome organization, such as the definition of hub proteins, but they typically lack the contextual information that relates specific assemblies to the availability of cellular resources and cellular functional state. In essence, the systematic exploration of the modularity of the proteome has yet to achieve the transition from deconceptualized maps to measurements in the context of cellular state. To bring about this transition several layers of data are required: (i) an accurate map of the assembly composition; (ii) information about its spatial organization, such as size, subunit stoichiometry and organization, and formation principles; (iii) a description of its quantitative temporal alteration as function of the cell state (e.g. signaling activation); (iv) its cellular coordinates and the resources that enable and constrain its formation; (v) a conceptual/computational framework to integrate structure- and systems-level information.

In this study, we combined and further developed orthogonal mass spectrometric (MS) approaches to address these challenges. They include analyses of a target assembly by different MS modalities; a new differential native separation method to study its disassembly dynamics; an in-depth, time-resolved determination of its absolute stoichiometry (AP-AQUA-MS); the combination of absolute quantification in lysates (lysate-AQUA) and affinity-purified samples to quantify the proportion of cytosolic proteins assembled in complex; and a framework to combine all these data layers in a model (Fig. 1). We selected as case study the tumor necrosis factor-receptor signaling complex (TNF-RSC), a signalosome that plays a pivotal role in both inflammation and cancer (*5,6*). This system was selected for three reasons: (i) it represents a challenging example of low-abundant, membrane-bound signalosome that forms only transiently and in a PTM-dependent fashion; (ii) there is a large body of literature that can be used to benchmark our results; (iii) in spite of (ii), several important questions remain unaddressed regarding its regulation, structural organization and systems-level properties. These include, for instance, regulation by phosphatases and architecture of the partaking protein complexes. In summary, we present a generic, widely applicable strategy for the contextual modelling of protein assemblies, and by applying it to a challenging membrane-bound immunological signalosome, we provide new insights into its biochemical, structural and systems-level organization.

**Figure 1.**
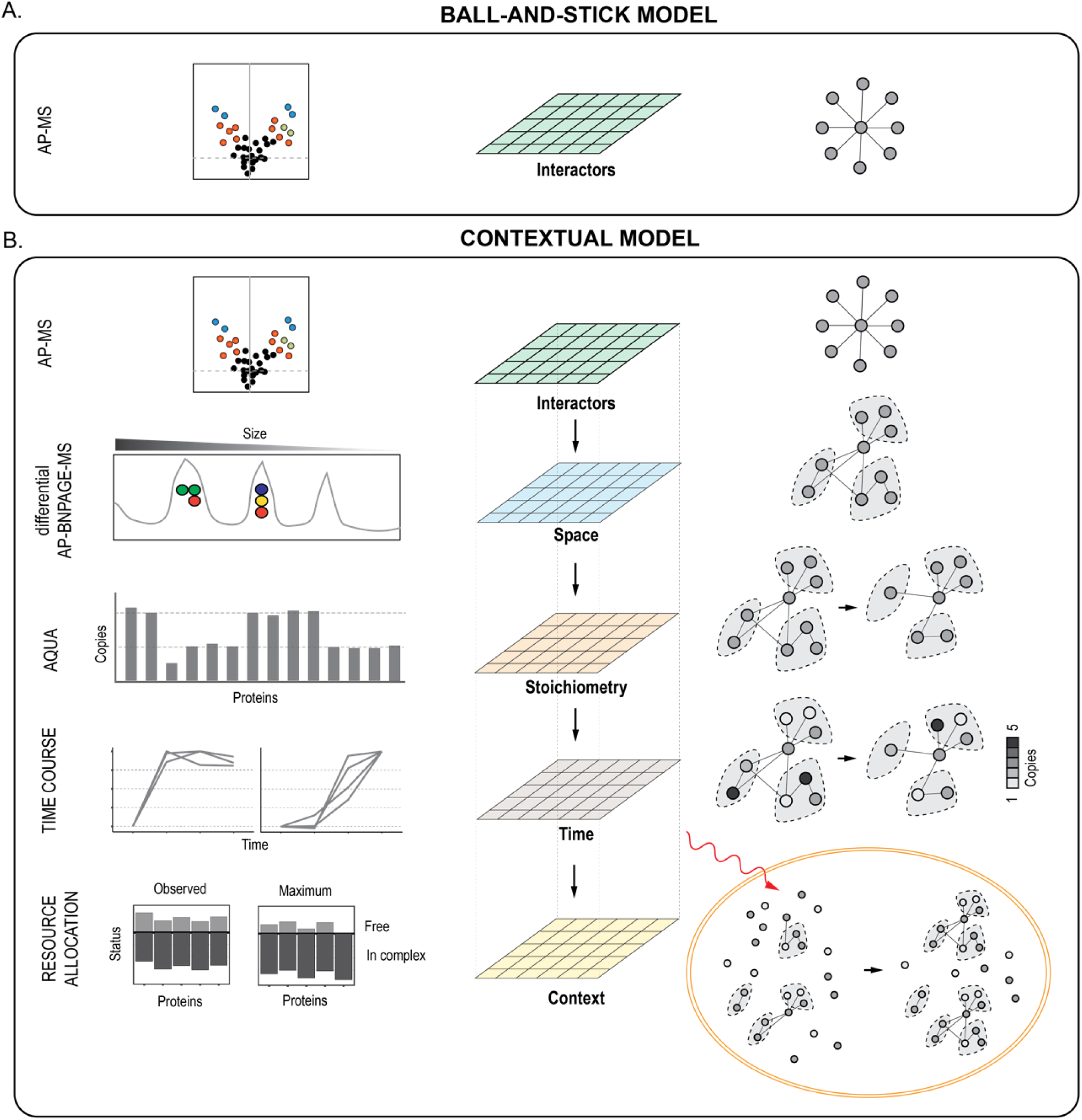
PPI contextual modelling approach. **(A)** In typical AP-MS workflow, PPI networks are generated from a single layer of data: those proteins that are identified as differentially abundant against a control (volcano plot, left). (**B)** To achieve a contextual modelling, we built on the first interaction layer, and further characterized a target assembly by a novel differential native separation by BNPAGE (AP-BNPAGE-MS), absolute quantification (AQUA) over time after stimulation, and determination of cellular resources distribution during signaling. The integration of these different layers can be used to describe constraints dictated by available resources and structural properties on the formation of assemblies and their activity.

## Results

### First layer of the contextual model: Identification of interactors using AP-MS (Fig. 2)

**Figure 2.**
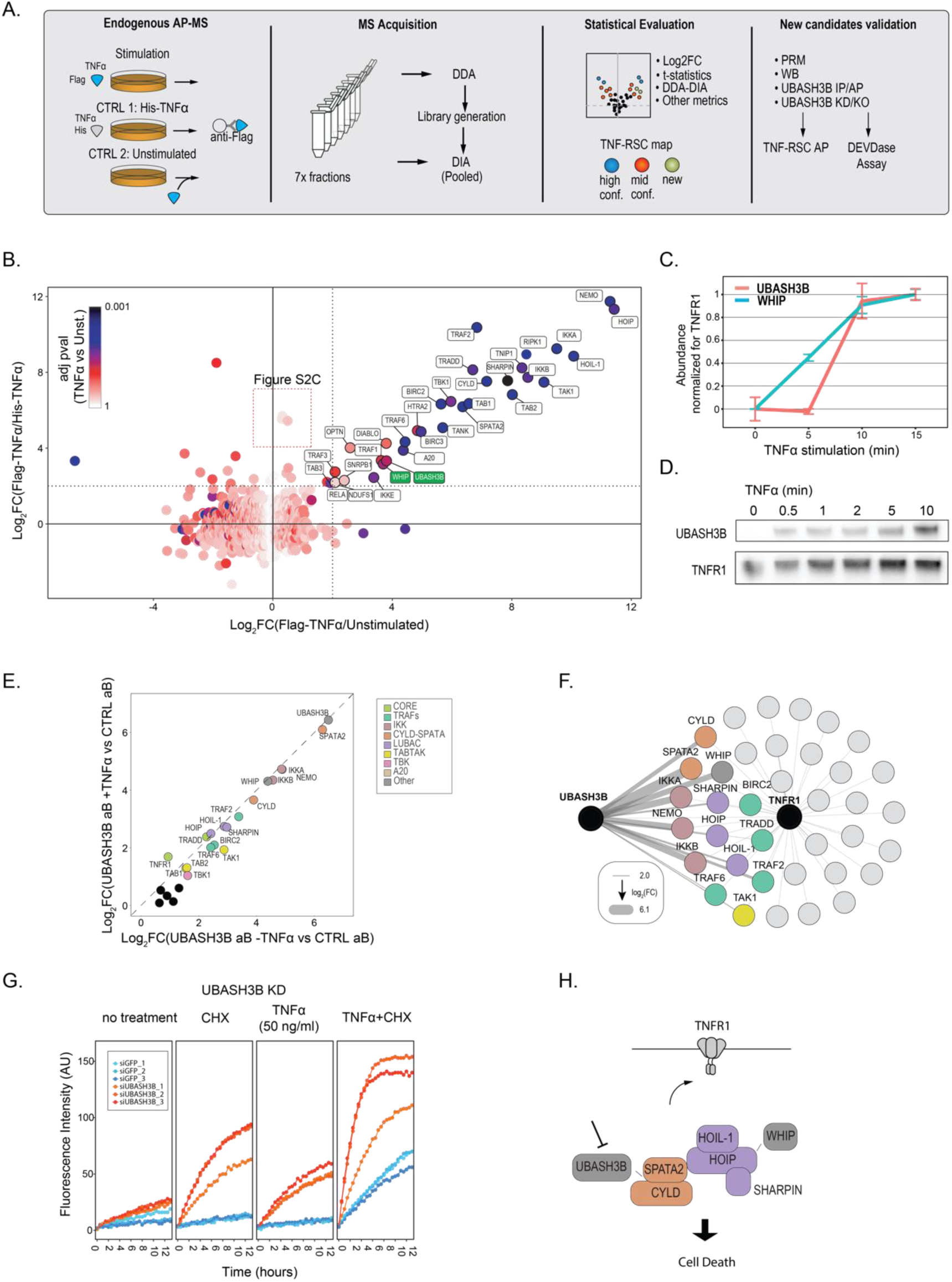
Landscape of TNF-RSC and characterization of UBASH3B as a new signalosome member. **(A)** Experiment design: A549 cells were stimulated and isolated using a Flag-tagged TNFα. Addition of Flag-tagged TNFα to unstimulated lysates or His-tagged TNFα to intact cells were used as controls. Samples were fractionated and analyzed by DDA, while pooled ones were analyzed by DIA. Standard statistical procedures were used to determine high confidence interactors, and additional analyses revealed mid-confidence associated proteins. Finally, additional pulldowns and biochemical experiments were performed to validate UBASH3B as a new TNF-RSC complex member. AP=affinity purification; IP=immuno-affinity purification. (**B)** Scatterplot showing protein enrichment across the two controls; adjusted *p*-value against the unstimulated control is coded in the dots color. (**C**) The recruitment of WHIP and UBASH3B to the TNF-RSC is confirmed by targeted proteomics on isolated signalosomes (A549 cells) across the indicated time points after stimulation. (**D**) Recruitment of UBASH3B to the TNF-RSC is further confirmed by immunoblot analysis. **(E)** Targeted IP-MS analysis of UBASH3B in A549 cells. The scatterplot displays the enrichment of the untreated and TNFα treated isolates against an unspecific serum control. (**F**) Network model showing overlap of proteins enriched with a log_2_FC threshold>2 in both UBASH3B IP-MS (left) and TNF-RSC affinity purification (right). Thickness of UBASH3B edges scales with enrichment of associated nodes. (**G)** DEVDase assay with the indicated siRNA (B10 from Table S12) against UBASH3B; siGFP was used as a control. Profile of each replicate is shown. (**H)** Model of the recruitment and signaling role of UBASH3B.

The composition of the TNF-RSC has been previously studied by MS following its isolation via tagged-TNFα (*7,8*). In the first step of our workflow, we set out to quantitatively map the TNF-RSC composition at a greater depth than previously reported. This was achieved by adding to the previous protocol a fractionation step, an orthogonal control (Material and Methods), and by adopting an analysis strategy based on multiple MS modalities (Fig. 2A). We first analyzed, by data dependent analysis (DDA) MS, TNF-RSC isolated from A549 cells stimulated for 10 minutes with TNFα (Fig. 2B, S1A/B/D/E/F; Table S1). As shown in the scatter plot in Fig. 2B, our results identify about 30 high-confidence interactors with a precision above 80% at log_2_FC>2, and thus recapitulate most of the previously known TNF-RSC interactors (Table S2). Remarkably, we identified two new high confidence (UBASH3B and WHIP, Fig. 2B,S2A) and several new or less well characterized TNF-RSC-associated proteins at a lower level of confidence. These include AZI2, TAX1BP1 (Fig. S2B/C) and KCTD2/5/17, which are enriched in a stimulus-independent fashion (Fig. 2B, inset; Fig. S2D; Supplementary Information for a more extensive discussion). One of the two novel high confidence interactors, UBASH3B, is a phosphatase known to regulate EGFR receptor internalization and TCR signaling (*9*). The other newly identified member, WHIP, is a triple ATPase. WHIP has been recently characterized as part of a trimeric complex involved in innate antiviral response (*10*). Furthermore, it also has been reported to interact with the ubiquitin ligase HOIP in a previous MS screen, but, to our knowledge, never been associated with the TNF-RSC (*11*). To confirm and characterize these interactions, we performed (i) data independent MS (DIA) analysis of the pooled fractions (Fig. S1C/G/H/I; S2E/F; Table S3); and (ii) targeted parallel reaction monitoring (PRM) analysis of UBASH3B and WHIP recruitment to the TNFR1 over time after stimulation (Fig. 2C). Both data sets overall supported our findings (Fig. 2C; S2E), which is further biochemically corroborated by immunoblot analysis (Fig. 2D, UBASH3B only). We next focused on UBASH3B. To understand its mechanism of recruitment to the TNF-RSC, we affinity-purified endogenous UBASH3B (A549 cell line; Fig. 2E; S3A-H; Table S4) and ectopically expressed UBASH3B (HEK293 cell line, C- or N-terminally tagged, Fig. S4A-F; Table S5) and quantified its interactors by MS. These experiments showed that (i) UBASH3B associates prior to receptor stimulation with TNF-RSC members (Fig. 2E; Fig.S4C); (ii) the association is transient or substoichiometric (based on absolute quantification estimates, as shown in Fig. S3B); (iii) that the interaction is likely mediated by CYLD-SPATA2 (Fig. 2E/F; S4C). Finally, we delineated the function of UBASH3B in the context of inflammatory signaling. To this aim, we first performed siRNA interference experiment in A549 cells and found that downregulation of UBASH3B by pooled (and one individual) siRNA leads to an increased apoptotic response, as measured by DEVDase assays with addition of cycloheximide and/or TNFα at different concentrations (Fig. 2G and S5A-C; Table S6). On the other hand, genetic ablation of UBASH3B in A549 cells does result in a mild pro-apoptotic response (Fig. S5D-F), and does not significantly perturb pro-inflammatory signaling, as measured by immunoblot analysis of phosphorylation of the downstream factors iκBα, p65 and MAP kinases (Fig. S6). Finally, we found that the composition of the TNF-RSC affinity purified from A549 UBASH3B knock-out (KO) cells does not significantly differ from the WT, suggesting that UBASH3B is not required for the signalosome stabilization or the recruitment of any of its analyzed members (Fig. S7 A-E; Table S7). Taken together, these data point at a potential, non-essential role of UBASH3B in regulating the apoptotic response (Fig. 2H). Overall, by using orthogonal MS approaches, and, critically, increasing the depth of our analysis, we could recapitulate prior knowledge about the TNF-RSC composition, identify new components of the signalosome and suggest the existence of a novel, UBASH3B-mediated layer of signaling regulation.

### Second layer of the contextual model: Spatial resolution using AP-BNPAGE-MS (Fig. 3)

**Figure 3.**
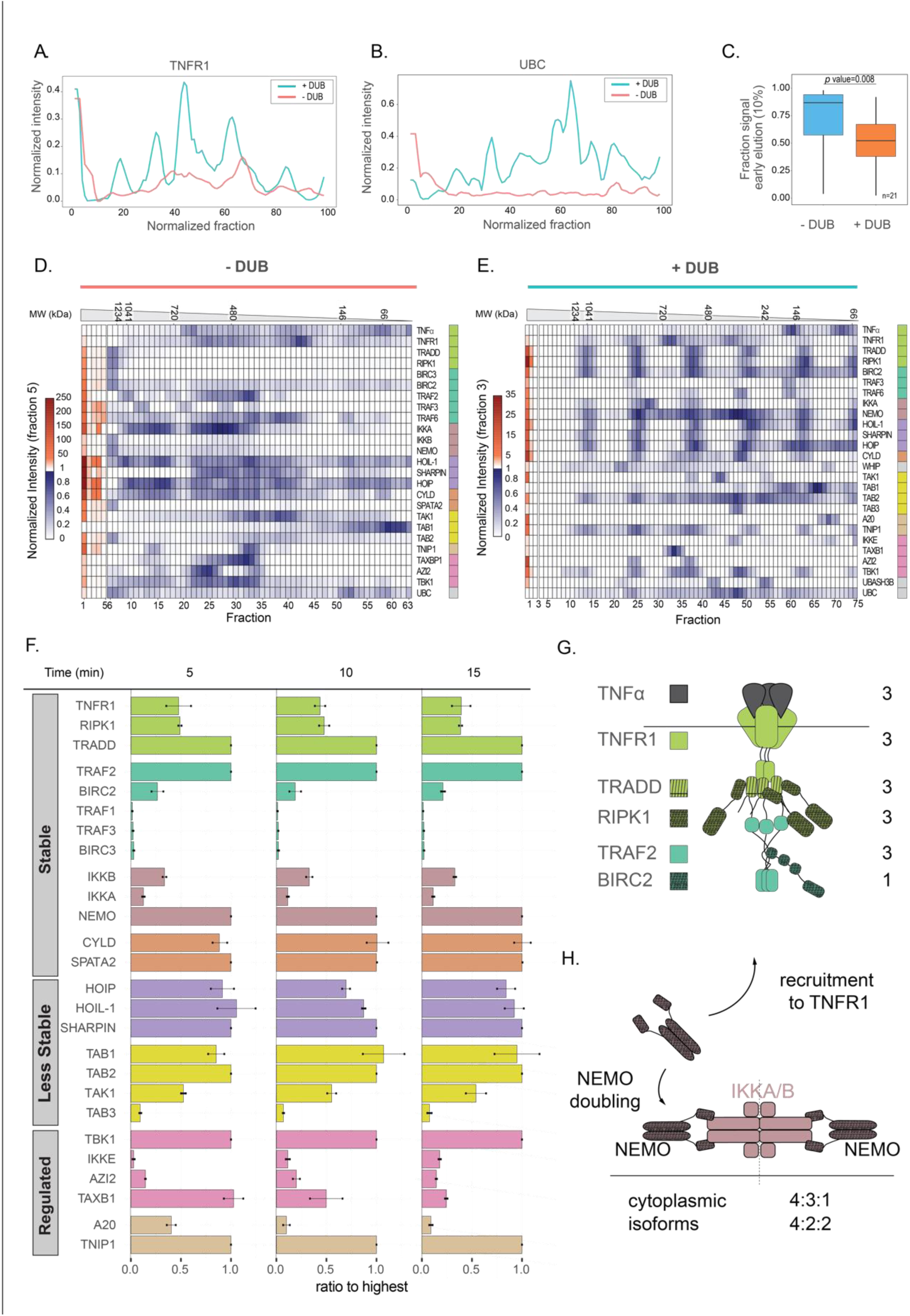
Architecture of the TNF-RSC. **(A/B)**. Comparison of TNFR1 receptor and ubiquitin signal in the two conditions (+/- DUB) indicates increase in signal and peak sharpening in DUB-treated complexes. Signal is normalized to the maximum intensity. (**C)** Signal is significantly shifted from early to late fractions in DUB-treated samples, indicating complex disassembly. (**D)** Signal distribution for the untreated, BNPAGE-separated TNF-RSC proteins. Signal normalized based on the intensity of fraction 5. (**E)** Signal distribution for the DUB-treated, BNPAGE-separated TNF-RSC proteins. Signal normalized based on the intensity of fraction 3. (**F)** Barplot of stoichiometries of complexes over time as determined by AP-AQUA-MS, and classified based on their stability. (**G)** Model of the TNFR1 core complex based on AP-MS (iBAQ) and AP-AQUA-MS data reflects results from previous *in vitro* characterizations. (**H)** Model of the stoichiometry rearrangement of the IKK complex upon recruitment, with duplication of the NEMO subunit and the existence of distinct cellular isoforms of the IKKA/IKKB complex.

We next aimed at defining the spatial organization of the signalosome and the complexes that constitute it. To this aim, we first developed a targeted, differential blue native polyacrylamide gel electrophoresis (BNPAGE) approach to evaluate the size and modularity of the TNF-RSC signalosome (Fig. S8A). In this approach, we (i) isolated endogenous TNF-RSC from a large amount of starting material (≥1×10^9 cells); (ii) separated the sample by BNPAGE prior to or after digestion with the generic DUB enzyme USP21 (Fig. S8B) (*12*), based on the rationale that the signalosome should be disassembled upon digestion of ubiquitin chains; (iii) used a precise and highly resolving gel slicing strategy, by which the BNPAGE gel is fractionated in 65-75 slices (Fig. S8C) and (iv) analyzed the protein contents of consecutive slices via the sensitive PRM method, providing sensitivity in the zeptomole range (1 zeptomol∼600 molecules; Fig. S8D-E, S9A-C; Table S8). We found that undigested TNF-RSC mostly remained trapped in the well, as evidenced by the stark accumulation of the signal in the first fractions of the gel, indicating that the isolation preserved the signalosome integrity and that its size exceeds the resolving power of the gel (∼1.5MDa; Fig. 3A/B, red trace; Fig. 3D; Fig. S10A). In contrast, USP21 digestion of the isolated TNF-RSC significantly improved its detectability (Fig. 3A-C, blue trace) and gave rise to a strikingly periodic signal distribution (Fig. 3E). This distribution indicated that several complexes, especially the membrane-proximal core components, the ubiquitin ligase complex LUBAC and the kinase complex IKK, occurred at regularly spaced intervals and with roughly constant intensity ratios across high molecular weight peaks (Fig. 3E; Fig. S10B/C). Besides recapitulating the ubiquitin chain-dependency of signalosome integrity and providing a lower boundary to estimate its size, these results point at a stoichiometrically regular arrangement of its constituting complexes.

### Third and fourth layers of the contextual model: Time-resolved stoichiometric measurements of the TNF-RSC following stimulation, using an AP-AQUA-MS approach (Fig. 3)

To infer accurate values for the stoichiometry of all the endogenous TNF-RSC complexes, we performed (i) AQUA-based absolute quantification of TNF-RSC over time post receptor stimulation (5,10,15 minutes); and combined it with (ii) iBAQ quantification of fractionated TNF-RSC, using 116 (AQUA) and 558 (iBAQ) peptides covering 30 proteins with up to 44 peptides/protein (Fig. S11 A-D; Table S9). Furthermore, (iii) we used as a benchmark i) a manually curated database of over 100 structural and biophysical publications (Fig. S11E; Table S10); and ii) re-evaluation of 3 proteome-wide SEC-MS datasets (*13, 14, 15*). On a general level, we found that AQUA and iBAQ-based values are in good agreement (Fig. S12 A-C), and that stoichiometries are remarkably stable over time post-stimulation, confirming in time what the AP-BNPAGE-MS had shown in space (Fig. 3F). Specifically, we grouped our findings in three classes: i) *Ex situ* validations of *in vitro* results; ii) Controversy-resolving findings; and iii) Novel findings (Supplementary Information for a more extensive discussion). Our data provides the first *in vivo* confirmation of the stoichiometry of the core signaling complex consisting of TNFR1, RIPK1, TRADD, TRAF2 and BIRC2/3. Based on iBAQ data, RIPK1, TRAF2 and TRADD occur in an approximate 1:1:1 ratio, or 3 copies each (Fig. 3G; Fig. S12D-F). AQUA measurements for the core complex indicated a different stoichiometry for TRADD, possibly due to specific biases in the selected TRADD peptides. Finally, BIRC2 was consistently found at an average ∼1:4 ratio with TRAF2, compared to a 1:3 ratio observed *in vitro* (*16*). Similarly, CYLD and SPATA2 are found to stably occur in a ∼1:1 relative stoichiometry, confirming that their extensive characterization *in vitro* reflects their *in vivo* organization (2:2 absolute stoichiometry)(*17*). The best fit for LUBAC is a (2 _HOIP_:2 _HOIL-1_:2_Sharpin_)+(1 _HOIL-1/Sharpin_) stoichiometry, where excess HOIL-1/Sharpin subunits are added to a core isostoichiometric complex. This is supported by crystallographic and biophysical evidence for an isostoichiometric arrangement of LUBAC (1:1:1 or 2:2:2)(*18,19*), by the existence of partial complex isoforms and by the potential dimerization of Sharpin via the SH domain (*20*).

The stoichiometry of the kinase complex IKK (NEMO:IKKA:IKKB) has been extensively investigated *in vitro* and dozens of potential solutions are compatible with the current data (*21, 22, 23, 24, 25, 26, 27, 28*). Our results, in combination with (a) absolute measurement of A549 cytosolic pools (Fig. S13A); (b) bioinformatic analysis of tissue-level data (Fig. S13B); (c) ratios derived from SEC-MS experiments (Fig. S12G); supports a model where the number of NEMO molecules in complex duplicates upon TNF-RSC recruitment, associating with two (or possibly more) distinct isoforms of the IKKA/IKKB (Fig. 3H). Importantly, this model reconciles and provides a rationale for several seemingly contradictory reports (Table S10; Supplementary Information). iii). Finally, our data indicates a 2_TAB1_:2_TAB2_:1_TAK_ ratio for the kinase complex TAB/TAK1 and an absolute stoichiometry of 2_TANK_:2_TBK1_ for the kinase TBK1 complex (Fig. S12D/E). In summary, by combining differential AP-BNPAGE-MS and stoichiometry measurements, we provide the first absolute quantitative, *ex situ*, time-resolved ensemble description of the TNF-RSC size, ubiquitin-regulation and complex stoichiometries.

### Fifth layer of the contextual model: Resource allocation of the TNF-RSC using the combination of AP-AQUA-MS and lysate-AQUA analysis (Fig. 4)

**Figure 4.**
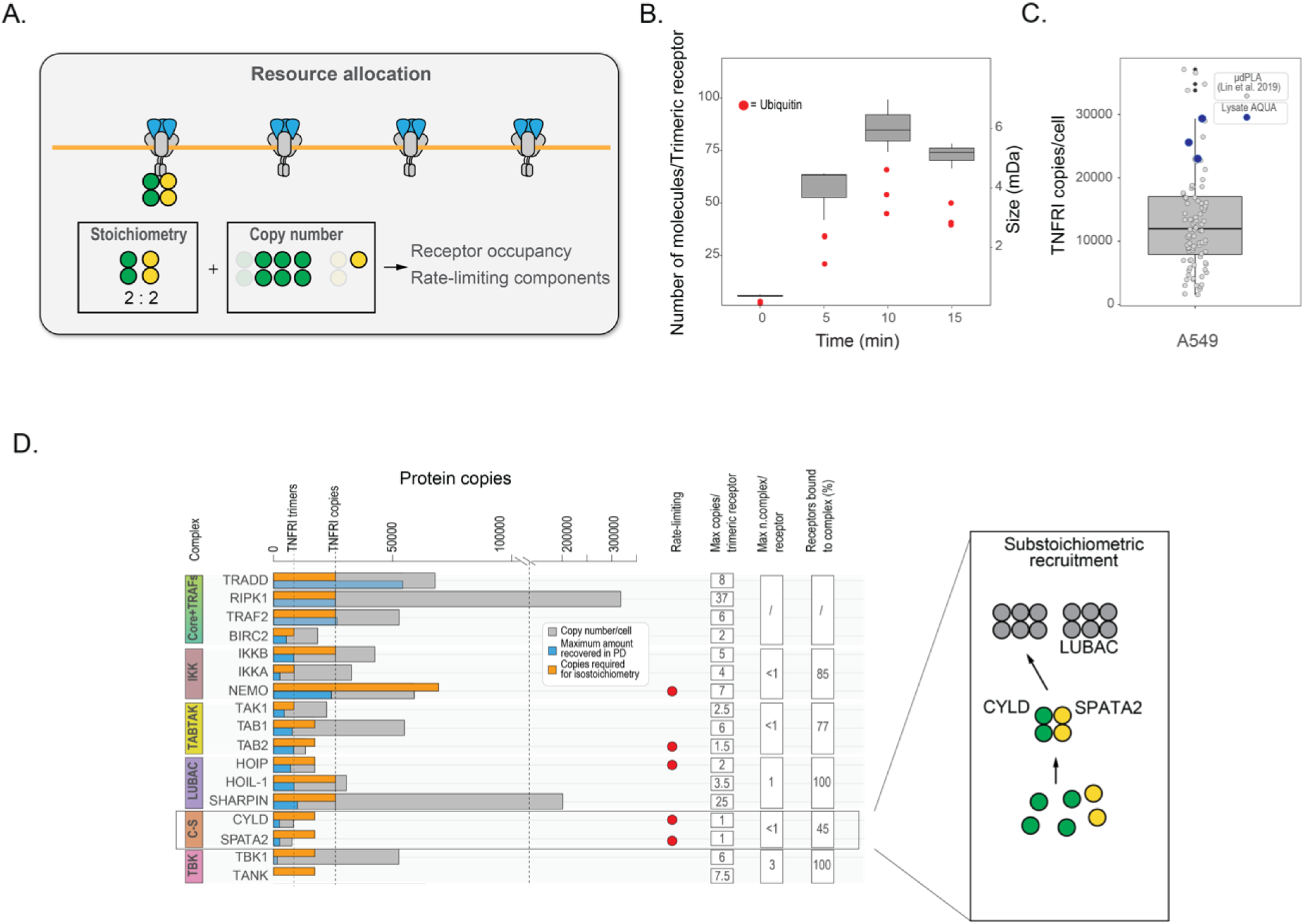
Cellular constraints on TNF-RSC formation. **(A)** Design: estimate of copy number/cell is combined with knowledge about stoichiometry to define receptor occupancy and identify rate-limiting complex-components. (**B)** MW and number of molecules of the TNF-RSC over time as estimated by AP-AQUA-MS. Red dots indicates estimated number of ubiquitin molecules. (**C)** Boxplot displaying lysate estimates of the number of molecules/cell of TNFR1 (blue dots), and the number determined in a nominally identical cell line by (Lin et al. 2019) using PLA (grey dots). (**D)** Resource allocation plot. Number of copies/cell (grey bars) are compared with estimated copies isolated from affinity-purified TNF-RSC (blue bars) and the number of copies required to achieve 1:1 stoichiometry with the receptor (orange bars), considering all stoichiometric relationships determined in Fig. 3F. For the deviation of TRADD from the maximum expected amount, see main text. (right) Combining knowledge about stoichiometry and cellular resources provides information about recruitment dynamics and of the CYLD-SPATA2 complex.

In the last step of our workflow we set out to model the abundance and formation of the TNF-RSC inside the cell. How many receptors are present on average in a cell? How many are going to form complexes? And what constrains their formation? To address these questions, we applied a simple modeling framework where information about TNF-RSC stoichiometries (as determined above) is combined with information about the protein copy number of the TNF-RSC members in the cell to determine several systems-level properties (Fig. 4A). They include (i) observed and (ii) maximum average molecular weight of the TNF-RSC; (iii) observed and (iv) maximum receptor occupancy (fraction of receptors forming a signalosome); (v) and rate-limiting components. We first calculated a lower boundary in the size of the TNF-RSC using the AP-AQUA-MS data, by estimating the average number of molecules (and their summed MW) associated with the receptor (Fig. 4B). We found that the signalosome size peaks at 10 minutes reaching a size of about 6MDa, a result that is compatible with the AP-BNPAGE-MS data, with ubiquitin making up the majority of its mass. We next estimated the protein copies/cell of all TNF-RSC proteins using 12 cell lysates collected before the AP-AQUA-MS experiment presented above (Fig. S13A; Table S11). We used the independent measurement of the protein copies/cell of the TNFR1 receptor as a calibrant, and benchmarked it against a recently published estimate carried out in A549 cells using an orthogonal and accurate method, proximity ligation assay (PLA) (Fig. 4C) (*29*). Next, to investigate how representative our cell line is, we compared our values with the deep proteome profiling of 29 human tissues (*30*) and found them to be overall in good agreement (r=0.769; Fig. S13B/C), and relatively stable around the mean of the proteome-wide abundance values (Fig. S13D). Third, to identify those proteins that are rate-limiting for the formation of the complex, we calculated the copies of proteins needed to form a complete signalosome given the stoichiometries determined in the previous section. This information is displayed in Figure 4D, where the cellular copy number of the TNF-RSC members (grey bars) is compared with the amount recovered from the affinity-purified signalosomes (blue bars) and the theoretical copy number needed to occupy all the receptors given the determined stoichiometries (orange bar). As expected, we found that proteins recovered in affinity purified samples never exceeded the estimated cytosolic amounts, and that, in general, the numbers are higher for receptor-proximal proteins (Fig. 4D, blue bars; with the exception of TRADD, see Materials and Methods). Moreover, our data identifies CYLD-SPATA2, NEMO and TAB2 as rate-limiting (Fig. 4D, red dot), and, on average, indicates that most complexes are present in less than one or at most one copy per receptor, even at the peak of recruitment (10 minutes). Finally, we asked whether this systems-level data allow us to make predictions about the composition of the signalosome and the effect of protein overexpression. To do this, we focused on CYLD-SPATA2, which is substoichiometric with respect to the TNFR1 receptor. Since we know that (i) CYLD and SPATA2 are bound in a 2:2 stoichiometry (Fig. 3F, Fig. S12D/E)(17), (ii) that endogenous CYLD and LUBAC components are more abundant than SPATA2 (Fig. S13 A/B), and (iii) that SPATA2 mediates CYLD recruitment to LUBAC (*7,17*), we would predict that only part of the cytoplasmic pool of CYLD is bound to SPATA2, and that as a consequence CYLD-SPATA2 complex is substoichiometrically bound to the LUBAC complex. This is indeed what our AP-MS shows: only ∼20-25% of LUBAC molecules are bound to CYLD-SPATA2 (Fig. S12D). Likewise, endogenous abundance of the many paralogs/redundant subunits present in the system (TRAF2/TRAF5; TRAF2/TRAF1; BIRC3/BIRC2; TAB2/TAB3; TBK1/IKKE) is enough to crudely predict their receptor-bound abundance, even though more detailed mechanistic knowledge is essential to understand their differential recruitment/activity mechanisms.

Overall, we present a framework to combine protein interaction data with estimates of absolute protein amounts to contextualize PPI inside the cell and define some of the constraints regulating their formation.

## Discussion

In this study, we apply and test the limits of an integrated MS-based interactomic pipeline, which mediates the transition from a generic, decontextualized identification of PPIs towards the generation of a quantitative PPI model in the context of specific cell states. The presented strategy alleviates several limitations of current proteomic approaches, specifically the sensitivity, quantification accuracy, and combination of data from different sources, and is based on the complementary contribution of several orthogonal techniques and acquisition methods. They each defined, layer by layer, the composition, spatial organization, stoichiometry, PTM- and signaling-dependence, as well as systems properties of a target assembly. As applied to the TNF-RSC, this workflow has provided novel glimpses into its composition, regulation, spatio-temporal organization and system-level constraints. With the first step of the workflow, we combined different MS acquisition modalities with sample fractionation and 2 orthogonal controls to map the TNF-RSC composition. We show that these simple incremental improvements can be critical in increasing the depth and confidence of identification, and capturing those interactions – substoichiometric or labile, but of potential relevance - otherwise buried in the noise. Of the several interesting candidates we have thus nominated, we carried out a characterization of UBASH3B, since phosphatases have been comparatively poorly characterized in the context of TNF signaling, and thus suggested a new layer of signaling regulation. In the second and third step, we resolve in space and time important architectural features of the TNF-RSC, information that can be readily incorporated in integrative structural models. The differential AP-BNPAGE-MS analysis provided a new dimension to understand the architecture of the TNF-RSC, and its results show that the determined interactome is organized in a highly modular fashion, as expected from the ubiquitin-dependency of its assembly. It is important to bear in mind that the analysis of this data remains at present confounded by several factors, including disassembly of the signalosome and limit of detection of low-abundant components, which can result in migration patterns not consistent with their known MW. To our knowledge, this is one of the first examples of an endogenous complex being separated on a fractionation dimension in combination with MS, but we expect that such experiments will be routinely incorporated in interactomic studies, as they will become increasingly less time-consuming and labor-intensive. On a similar vein, there are only few publications that have used bottom-up MS to decode the absolute stoichiometry of specific complexes (*31,32*). In spite of the challenges entailed, we show that stoichiometry determination on pulldown samples represents a unique vantage point to capture ensemble, contextual structural information locally (pulldowns) and globally (lysate). This information is essential to interpret and validate *in vitro* data, but also to provide novel, self-contained and robust insights. Finally, on a systems level, at least two seminal studies have examined the role of protein abundance in other signaling systems (*33,34*), but have not taken into account the specific structural and stoichiometric requirements that proteins need to meet in order to be functional. We show here that by combining knowledge about stoichiometry, absolute abundances in the isolated signalosome and lysates, we can define lower and upper boundaries in the composition and formation of the signalosome in question, and make educated predictions on recruitment stoichiometries. To conclude, one of the greatest challenges of current biology is the modeling of the proteome in context, and interactomic studies have much to contribute to this endeavor. However, their use will remain limited unless they will incorporate quantitative, context- and structure-informed aspects. Our study describes a platform to achieve such integration.

## Supplementary Figures

**Figure S1.**
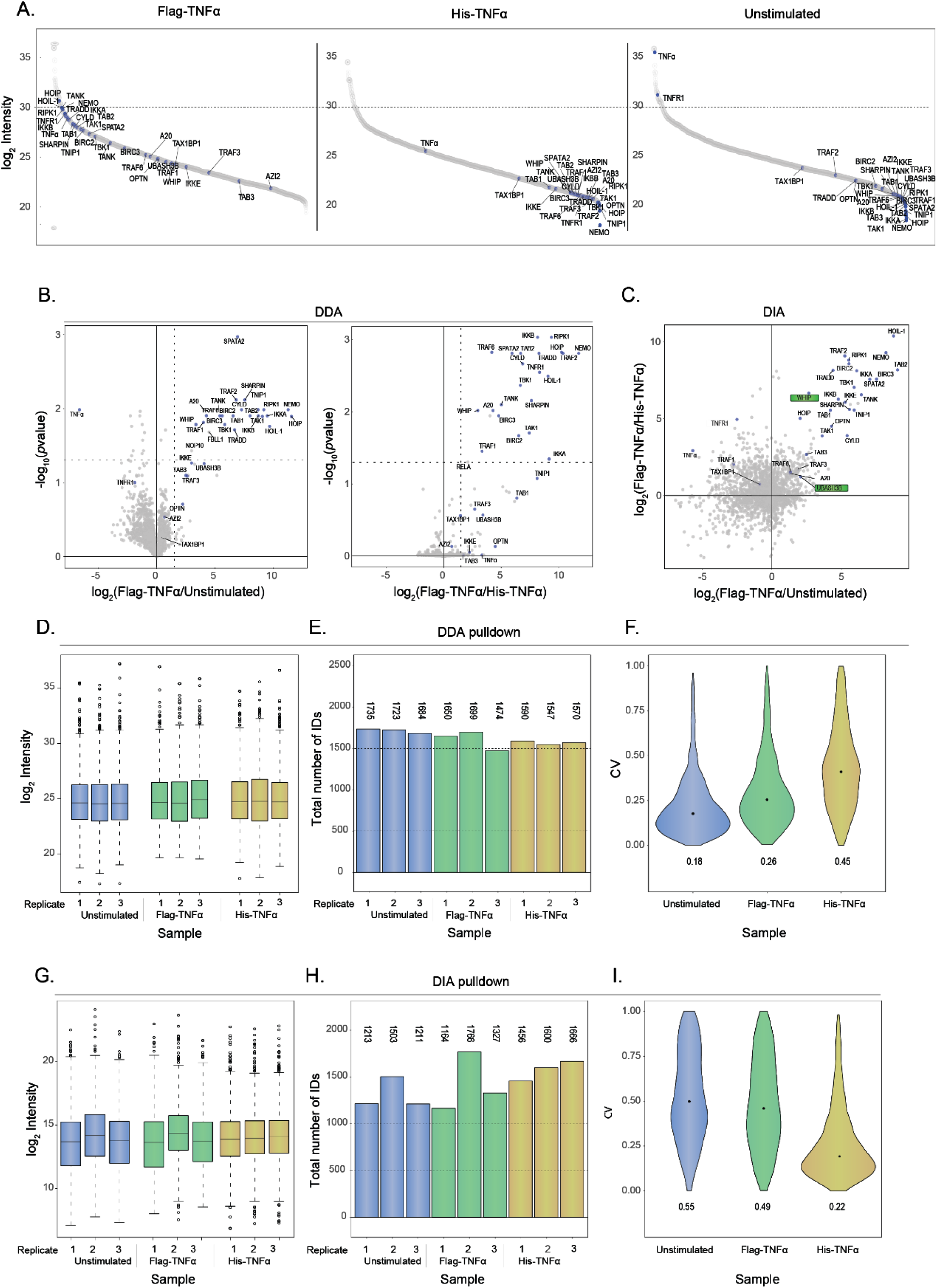
Quality controls on affinity-purified TNF-RSC analyzed by DDA and DIA. **(A)** Intensity distribution (log_2_) of TNF-RSC members across treated and control samples acquired by DDA. (**B**) Volcano plots of individual controls (DDA data). (**C**) Scatterplot of log_2_FC values showing protein enrichment across two controls (DIA data; analogous to Fig. 2B). (**D/G)** Boxplot of log_2_ intensities distribution across samples (DDA and DIA data, respectively). (**E/H)** Number of protein IDs identified in individual replicates (DDA and DIA data, respectively). (**F/I)** Violin plot showing coefficient of variation values of raw data (DDA and DIA data, respectively).

**Figure S2.**
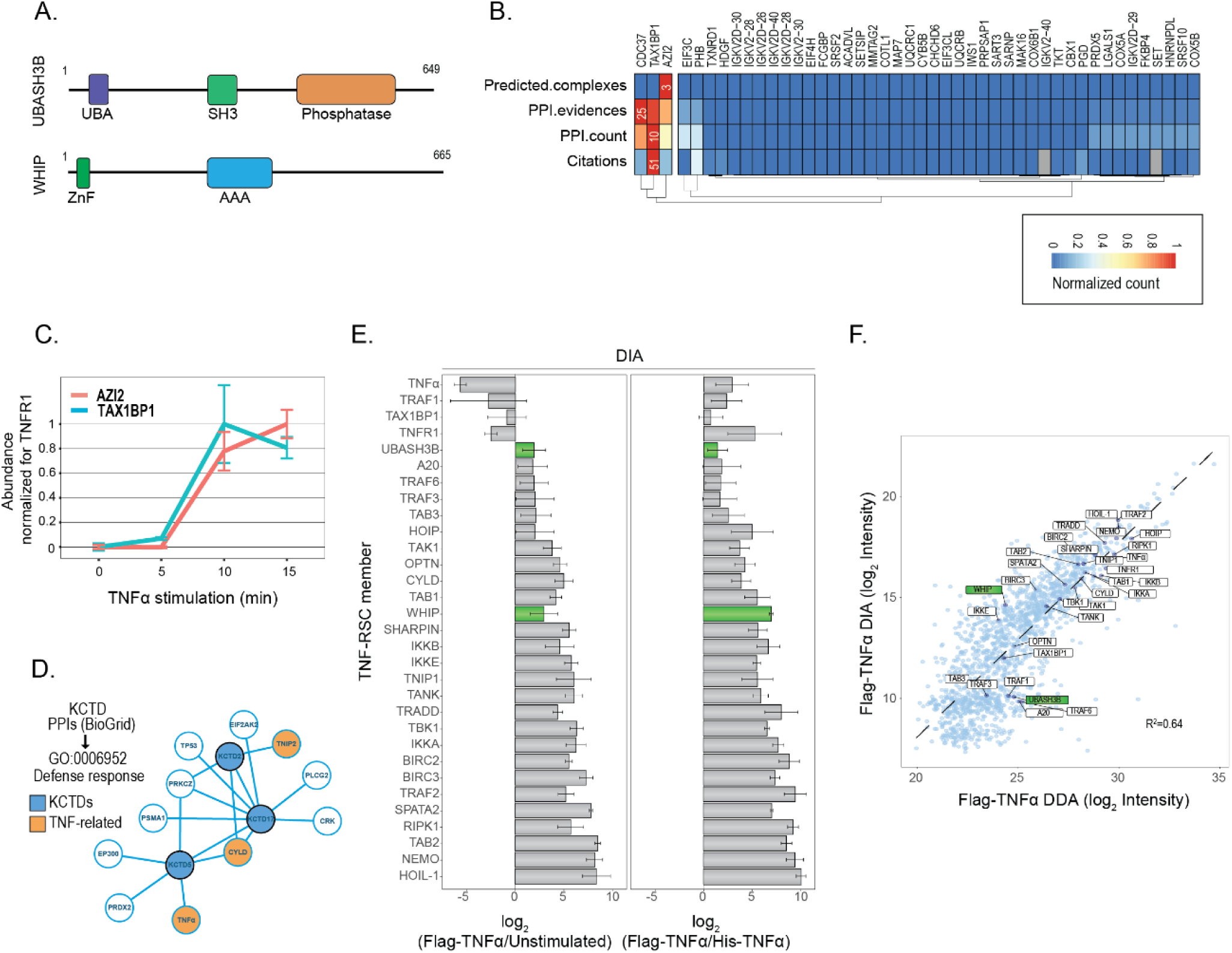
Identification of mid-confidence interactors and further validation by DIA of UBASH3B and WHIP recruitment to the TNF-RSC. **(A)** Primary sequence of UBASH3B and WHIP. (**B)** Clustering of mid-confidence interactors by multiple criteria isolates three known TNF-RSC associated proteins. (**C)** The recruitment of AZI2 and TAX1BP1 to the TNF-RSC is confirmed by targeted proteomics on isolated signalosomes (A549 cells) across the indicated time points after stimulation. (**D)** Previously reported interactions (Biogrid) of KCTD proteins indicate putative association with TNF-RSC members. (**E)** Log_2_FC enrichment of signalosome proteins identified in the treated and control samples from the DIA dataset. WHIP and UBASH3B are highlighted in green. (**F)** Scatterplot indicates correlation of protein intensities (log_2_) between DIA and DDA data. WHIP and UBASH3B are highlighted in green.

**Figure S3.**
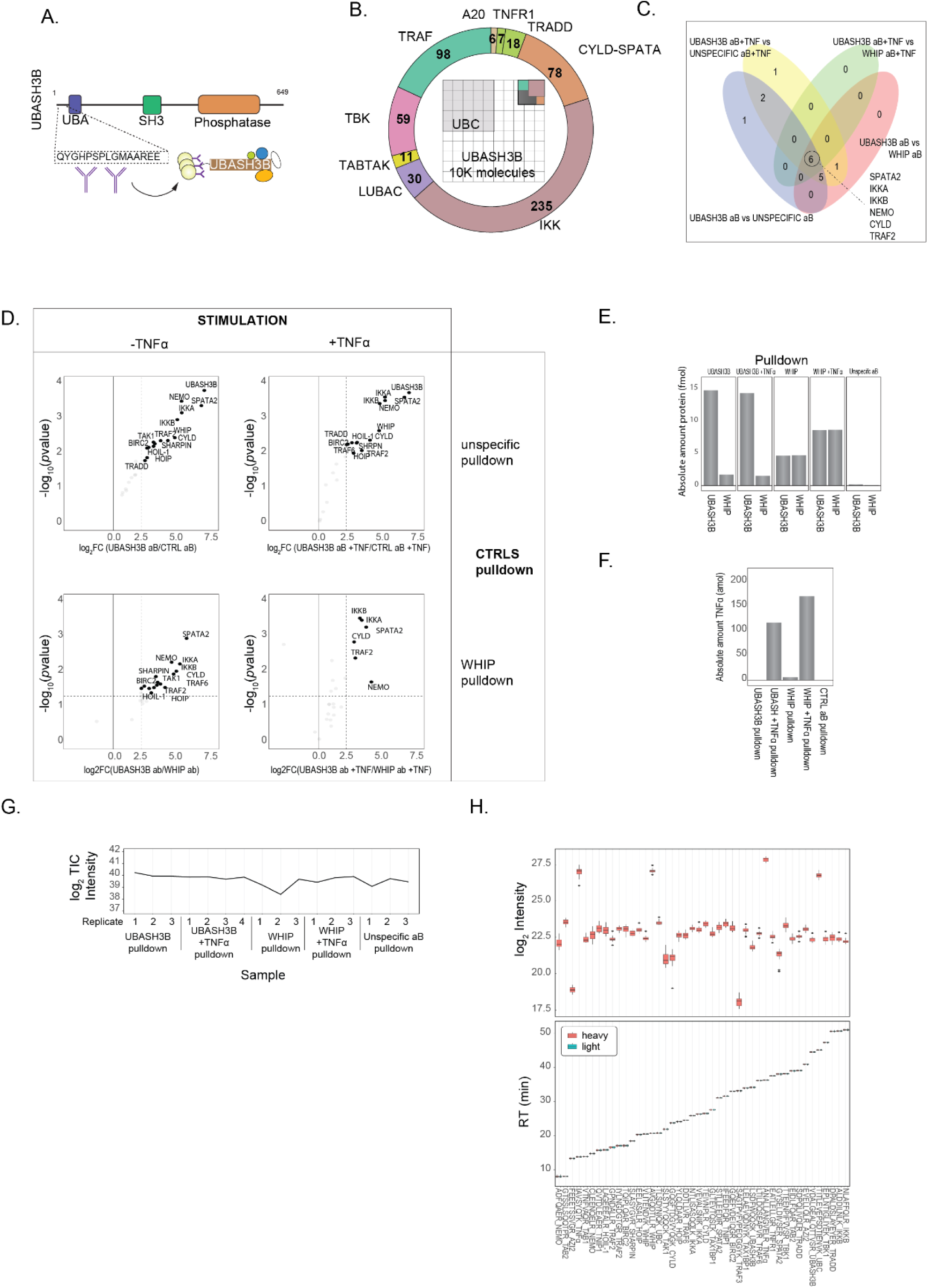
Characterization of the UBASH3B interactome by IP-MS in A549 cells. **(A)** Primary sequence of UBASH3B and peptide used to raise antibody for endogenous IP-MS. **(B**) Occupancy plot (inside) showing the fraction of the indicated interactors bound to an arbitrary number of UBASH3B molecules; the numbers are reported in the circular doughnut chart. (**C)** Venn diagram with proteins identified as consistently significant across the four conditions shown in panel S3D. Four conditions correspond to UBASH3B IP-MS against IP-MS with unspecific antibodies (serum control) +/-ligand and WHIP IP-MS +/-ligand. The WHIP control was used as an additional, more stringent control. Since UBASH3B and WHIP antibodies were raised together and purified sequentially, this control also addresses concerns regarding potential antibodies cross-contamination. (**D)** Volcano plots of UBASH3B endogenous affinity purification against four controls, as described in panel C legend. (**E)** Average amount of WHIP in UBASH3B in pulldowns described in panel C. (**F**) Average amount of ligand in the indicated conditions. (**G)** Total ion current (TIC) from UBASH3B IP-MS targeted analysis. **(H)** Signal from reference heavy peptides (top) and consistent pairing of heavy and light peptides (bottom) measured in UBASH3B IP-MS sample.

**Figure S4.**
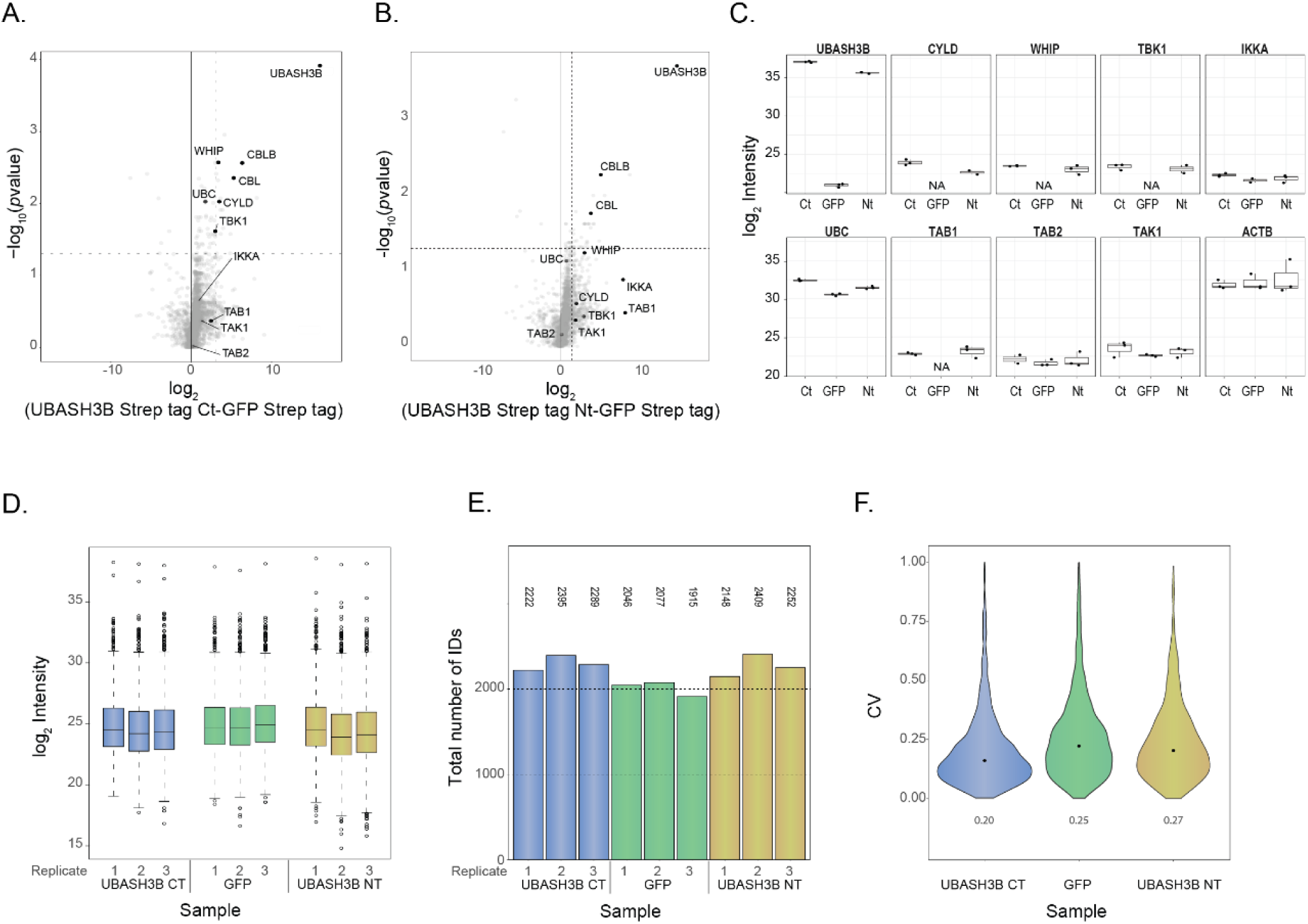
Characterization of the interactome of ectopically expressed UBASH3B by AP-MS in HEK293 cells. (**A)** Volcano plot of affinity purified C-terminally tagged UBASH3B against a GFP control indicates significant enrichment of TNF-RSC proteins. (**B**) Volcano plot of affinity purified N-terminally tagged UBASH3B against a GFP control indicates enrichment of TNF-RSC proteins. (**C)** Abundance of selected TNF-RSC proteins in AP-MS of C- and N-terminally tagged UBASH3B as well as GFP analyzed by DDA. Actin is shown as control. (**D)** Boxplot of log_2_ intensities distribution across samples. (**E)** Number of protein IDs identified in individual replicates. (**F)** Violin plot showing coefficient of variation values of raw data.

**Figure S5.**
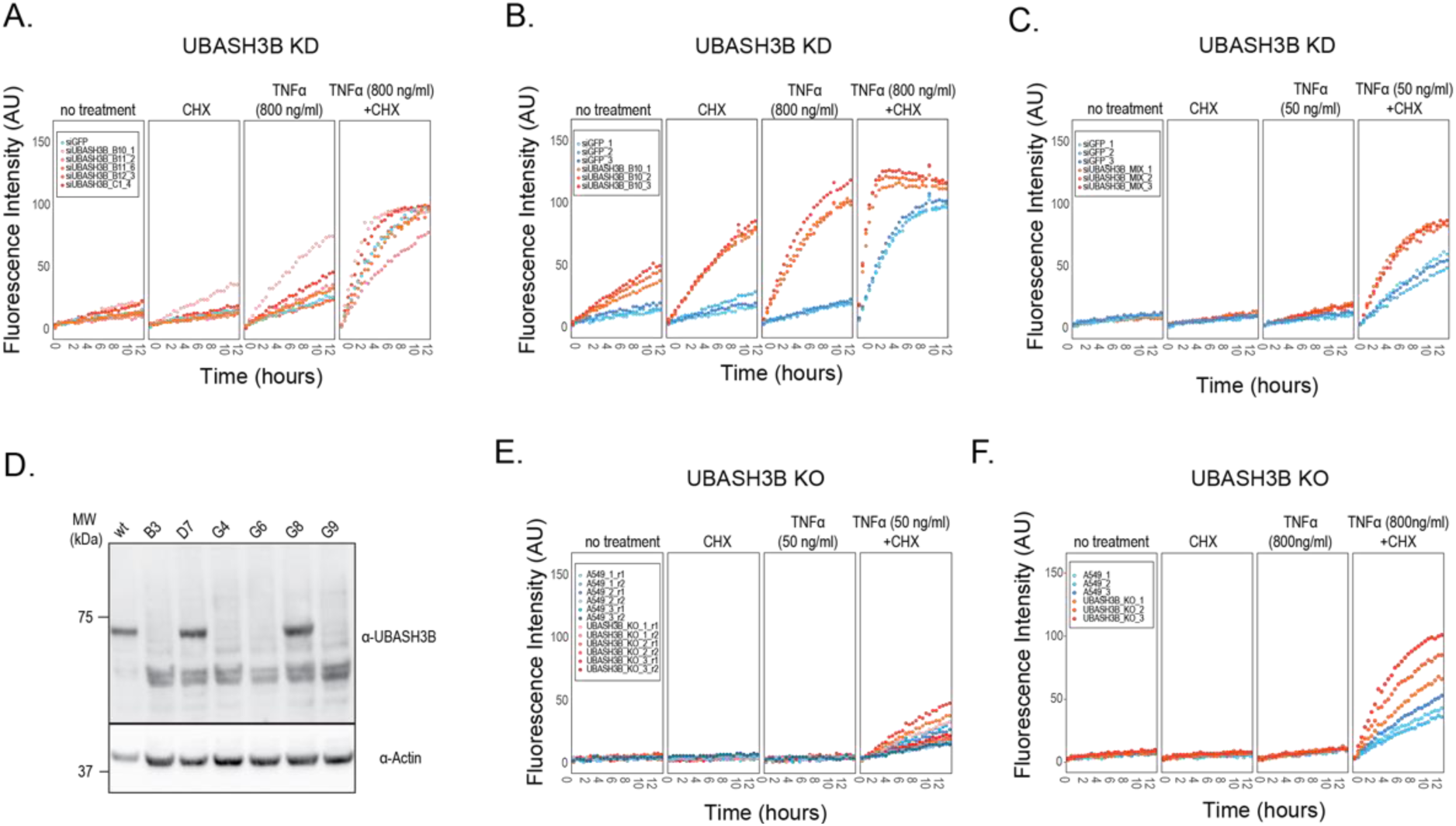
Apoptosis assay in UBASH3B KD (knock-down) and KO A549 cells. **(A/B/C)** DEVDase assay with the indicated siRNAs and ligand concentrations in UBASH3B KD A549 cell line. (**D**). Immunoblot against UBASH3B in clones of UBASH3B KO A549 cells. (**E/F**) DEVDase assay with the ligand concentrations in UBASH3B KO A549 cell line.

**Figure S6.**
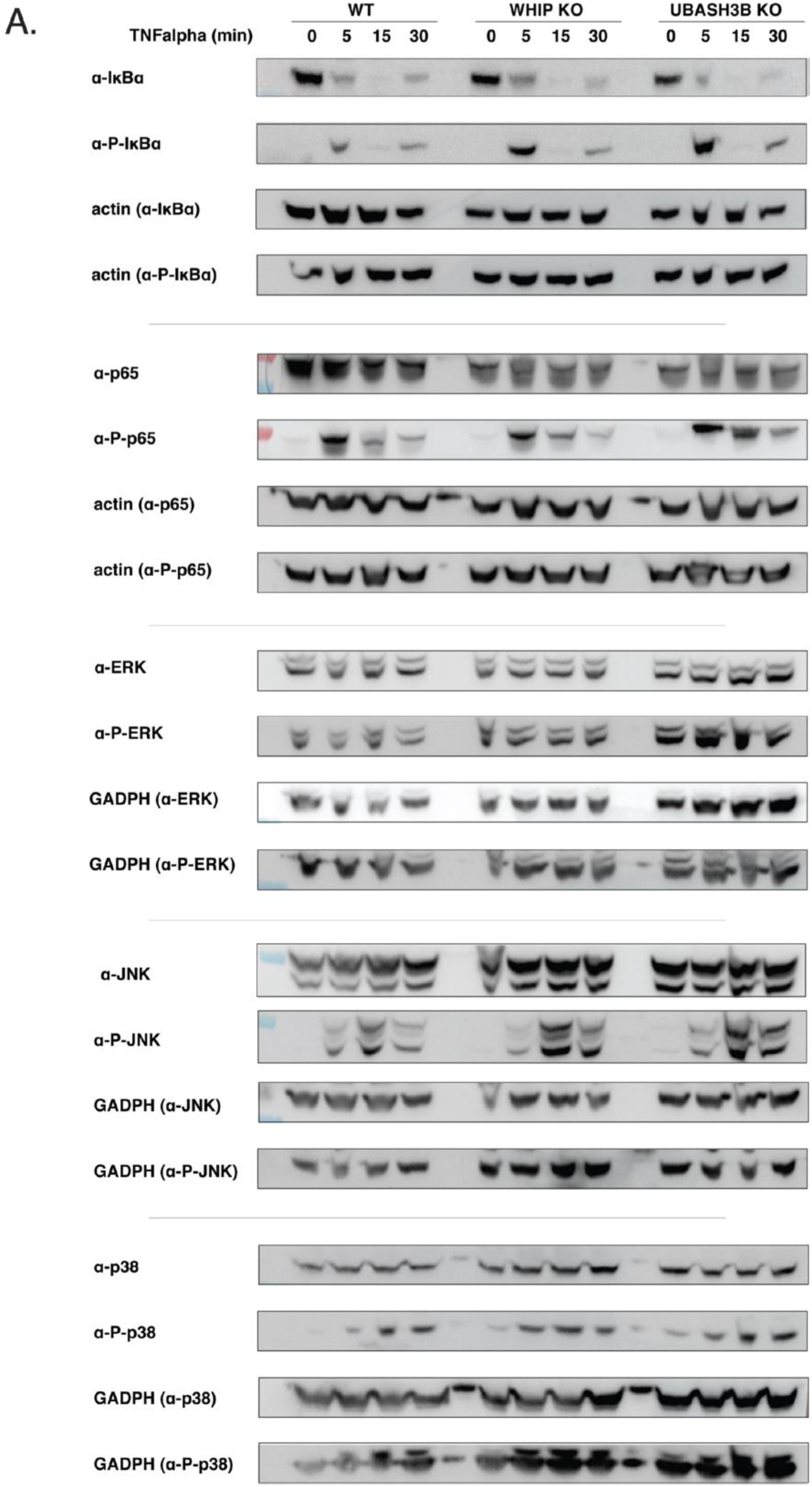
Immunoblot analysis of signaling progression in UBASH3B KO A549 cell line. Immunoblot against the indicated proteins in A549 WT/UBASH3B KO/WHIP KO across the indicated time points after TNFα stimulation.

**Figure S7.**
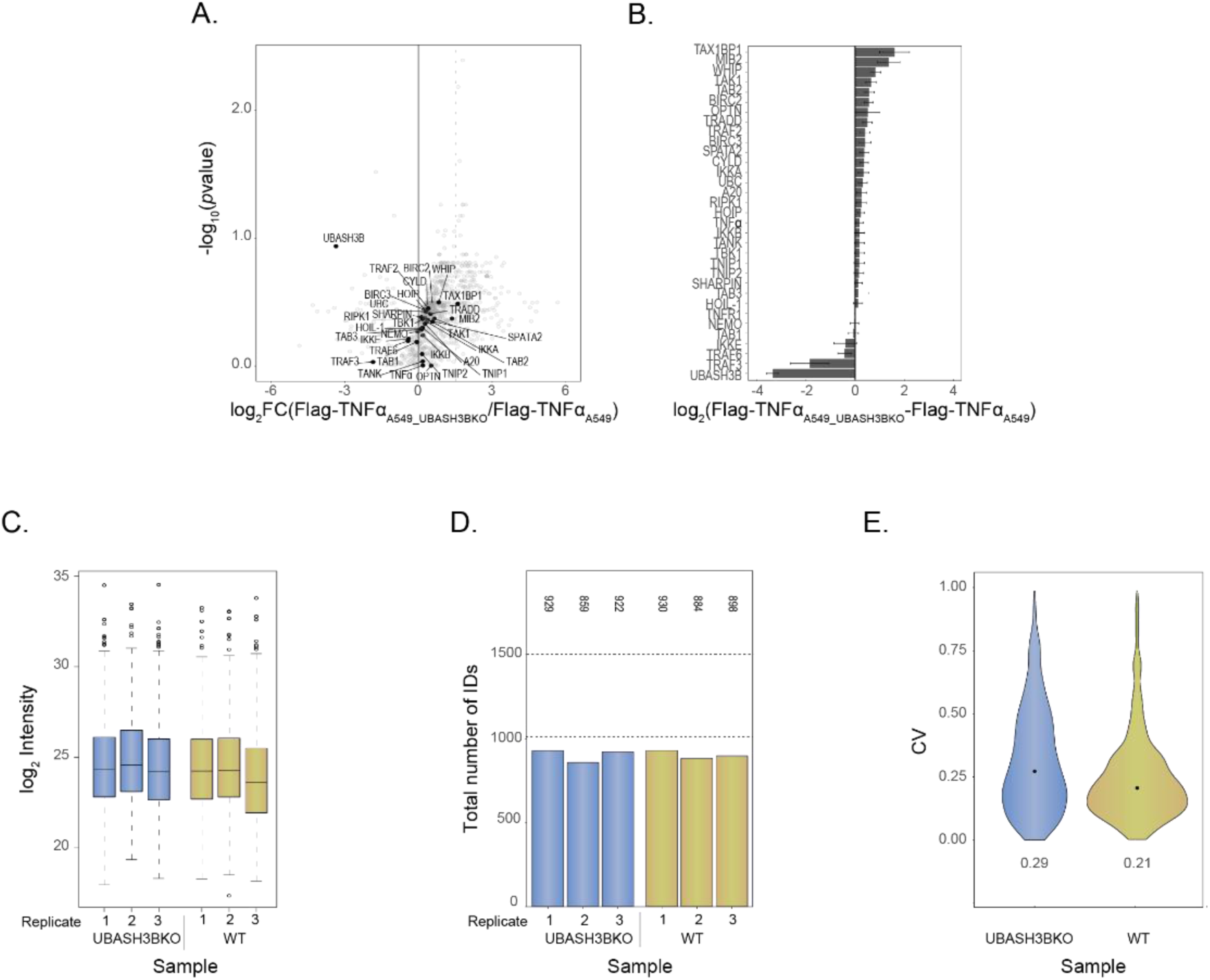
Characterization of the TNF-RSC interactome in UBASH3B KO A549 cells. (**A)** Volcano plot of TNF-RSC affinity purified with tagged TNFα in UBASH3B KO A549 cells against using WT A549 cells as a control. (**B**) Log_2_FC enrichment for selected TNF-RSC members. (**D)** Boxplot of log_2_ intensities distribution across samples. (**E)** Number of protein IDs identified in individual replicates. (**F)** Violin plot showing coefficient of variation values of raw data.

**Figure S8.**
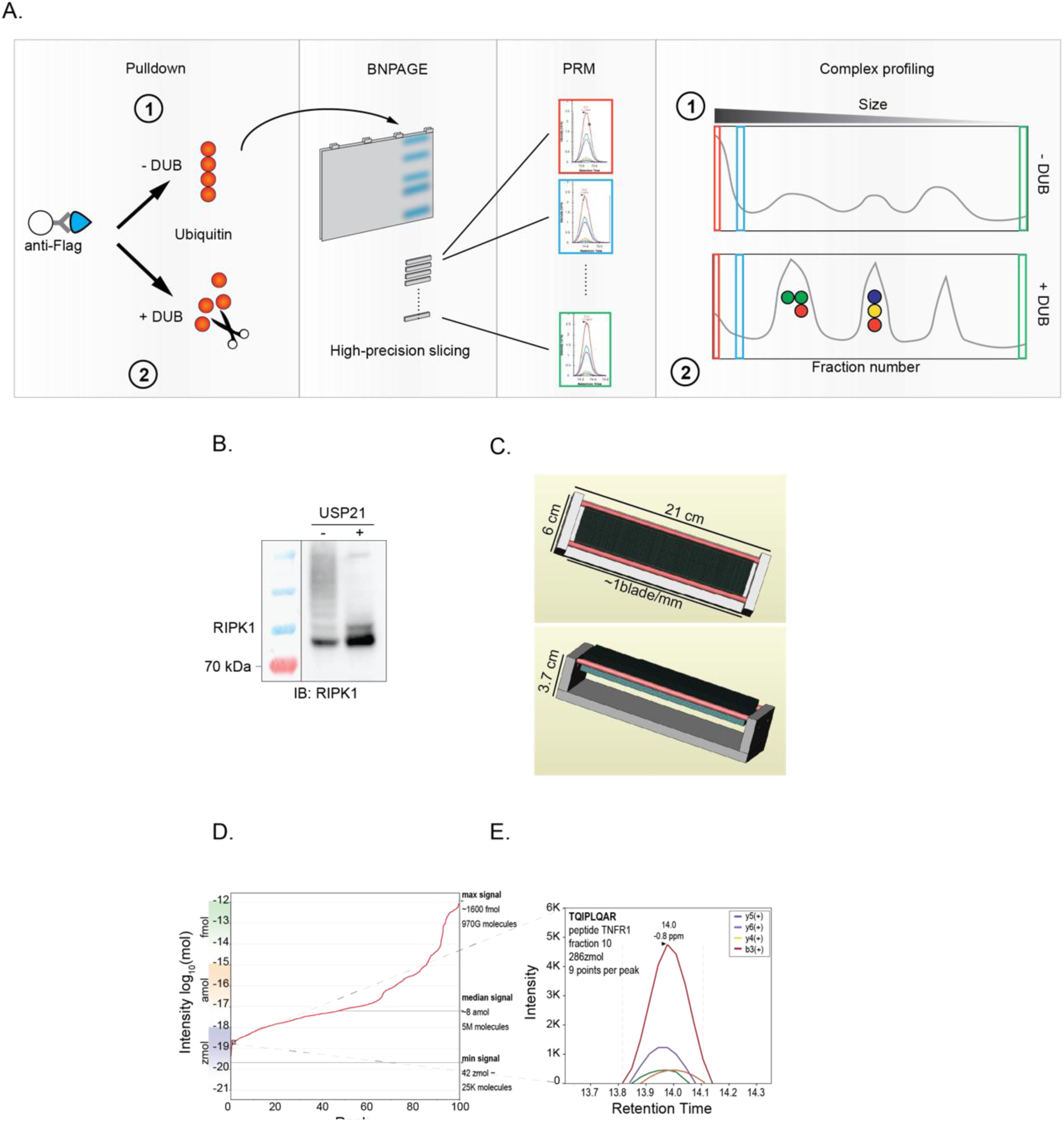
Experimental design of the TNF-RSC fractionation by BNPAGE. (**A)** Experiment design: affinity purified TNF-RSC with or without prior addition of the DUB USP21 were loaded and separated onto a BNPAGE gel. The gel was sliced with a custom-made gel slicer (panel **C**) and the extracted, digested proteins were analyzed by targeted proteomics. (**B)** Treatment of affinity purified TNF-RSC with USP21 induces a nearly quantitative reduction of ubiquitin chain signal as assessed by RIPK1 immunoblot. (**C**) Rendering of the high-precision gel slicer custom-designed and -produced (ETH Physics workshop) and used in this study. (**D)** Distribution of peptide amounts as estimated from average heavy AQUA standards run at regular intervals between fractions (USP21-treated sample). While most signal falls within the attomole range, limit of detection of peptide is in the high zeptomole range. (**E)** Selected peak group from a peptide with an estimated abundance in the zeptomole range where not only detection but also reliable quantification was possible.

**Figure S9.**
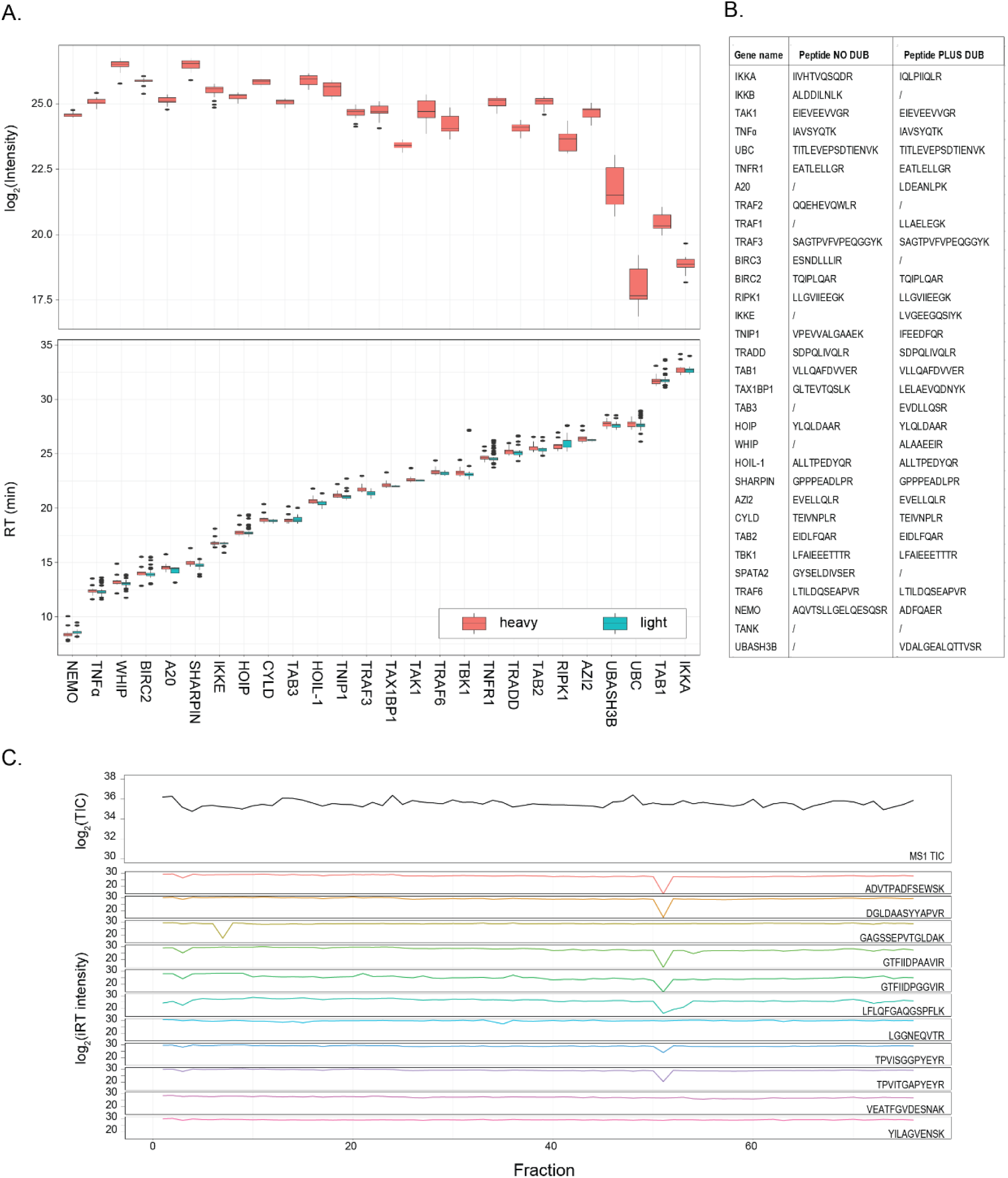
Quality controls related to the BNPAGE experiment. **(A)** Signal from reference heavy peptides (top) and consistent pairing of heavy and light peptides (bottom) measured in BNPAGE sample (treated with DUB USP21). (**B)** List of peptides monitored by PRM in the AP-BNPAGE-MS experiments. (**C)** Total ion current (TIC) profile (top) and intensity of iRT peptides (bottom) from the sample treated with DUB USP21.

**Figure S10.**
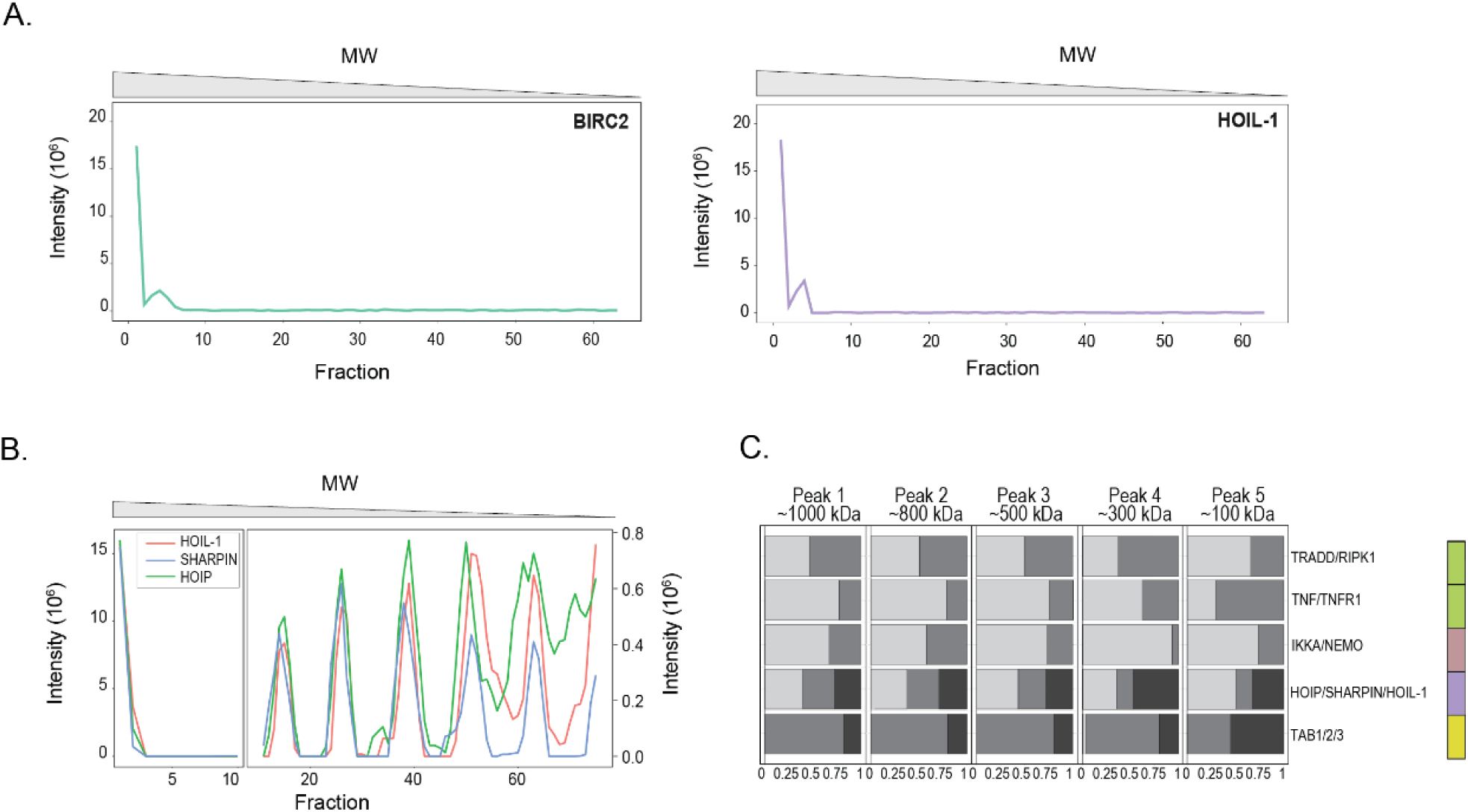
AP-BNPAGE-MS profiles of TNF-RSC untreated or treated with DUB. (**A)** Signal distribution for selected proteins (BIRC2 and HOIL-1) reveals protein accumulation in the stacking region of the gel. (**B)** BNPAGE migration profiles of LUBAC members after treatment with DUB USP21. (**C)** Relative abundance of membrane proximal proteins (green), IKK (brown), LUBAC (purple) and TAB/TAK complex members (yellow) remain constant across the first three peaks of the BNPAGE after treatment with DUB USP21.

**Figure S11.**
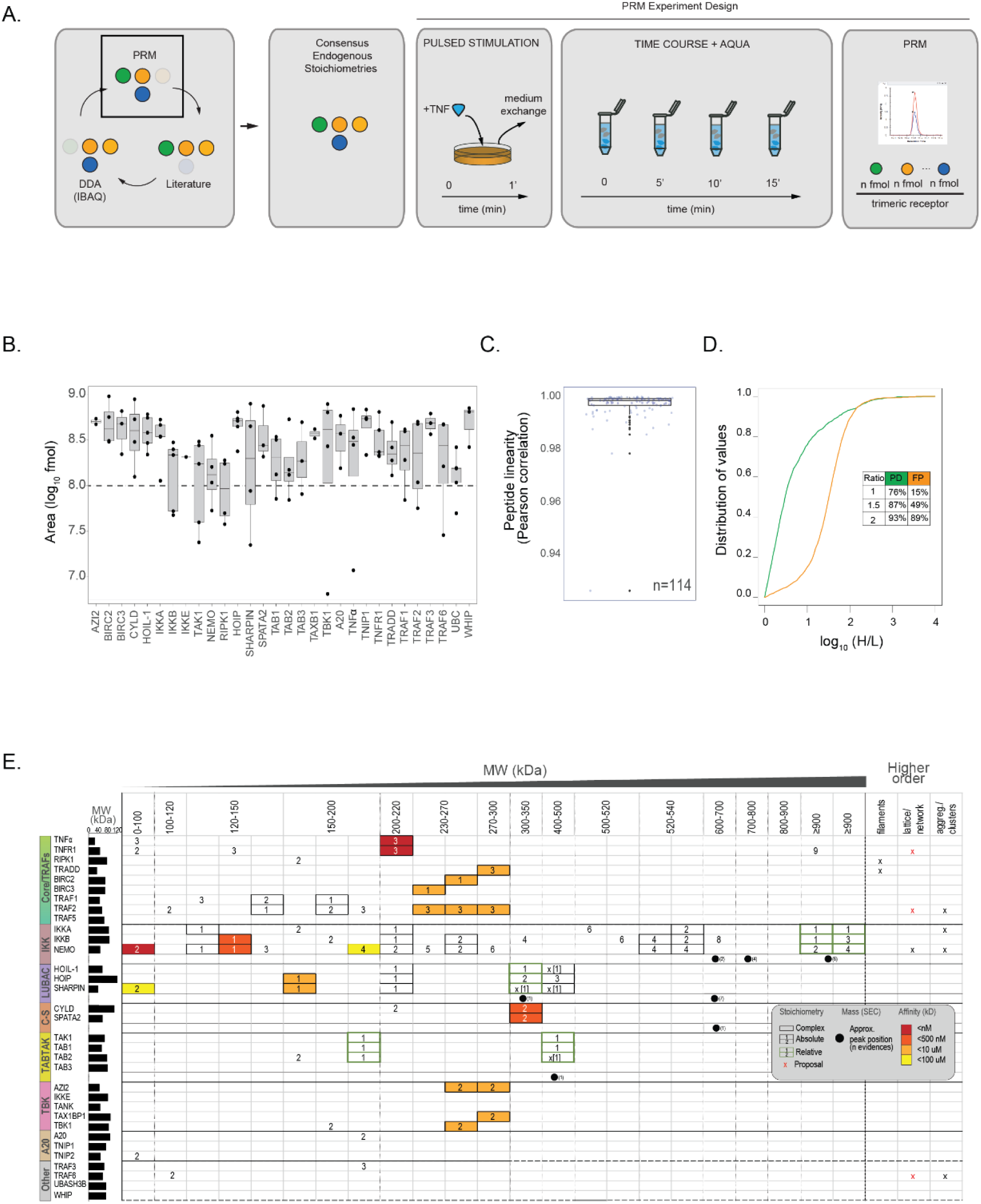
Design, quality controls and benchmark of AP-AQUA-MS experiment. **(A)** Design (left): Stoichiometries obtained by DDA and AQUA were compared with values from the literature and a consensus stoichiometry for each of the complexes proposed. AQUA experiment design (right): A549 cells were pulse-stimulated with TNFα and samples collected at 4 time points (time 0 was used as a control). AQUA peptides for TNF-RSC proteins were added to the samples and quantified by targeted proteomics. (**B**) Intensity of the AQUA peptides used in this study falls for about 80% of the peptides within 1 fold log_10_ intensity. (**C)** Peptides exhibit linear behavior over up to six orders of magnitude. (**D)** Intensity difference between heavy and light peptides used to quantify the affinity purified TNF-RSC and the proteins present in the lysate. (**E)** Summary of the literature curation, representing possible homooligomeric and heterooligomeric states for each TNF-RSC protein and complex (y axis) across the theoretical or observed molecular weight range (x axis). Selected data about MW as observed by SEC-WB and affinity of protein-protein interactions is also shown.

**Figure S12.**
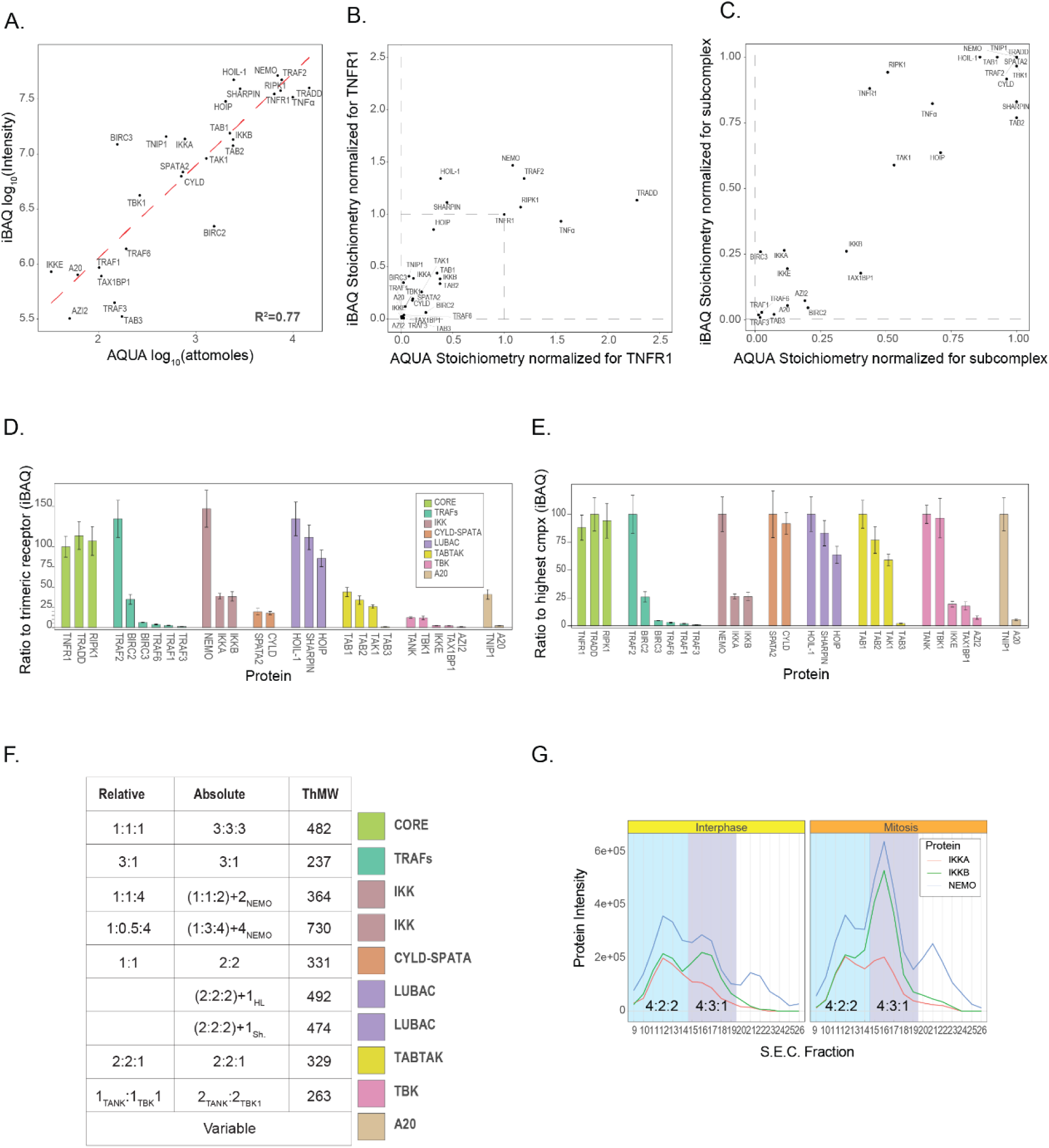
Assessment of TNF-RSC complexes stoichiometry by orthogonal approaches. **(A/B/C)** Correlation between stoichiometries estimated by AP-AQUA-MS and AP-MS (iBAQ) using log_10_ intensities (**A**), stoichiometries normalized to the receptor TNFR1 (**B**) and to the most abundant complex member (**C**). (**D)** iBAQ quantification of TNF-RSC AP-MS DDA data normalized to the trimeric TNFR1 receptor. (**E**) iBAQ quantification of TNF-RSC AP-MS DDA data normalized to most abundant complex member. (**F)** Table reporting suggested relative and absolute stoichiometries for the indicated complexes. (**G**) Reanalysis of data from (Heusel et al., 2020) indicates the existence of two cytoplasmic isoforms of the IKK complex.

**Figure S13.**
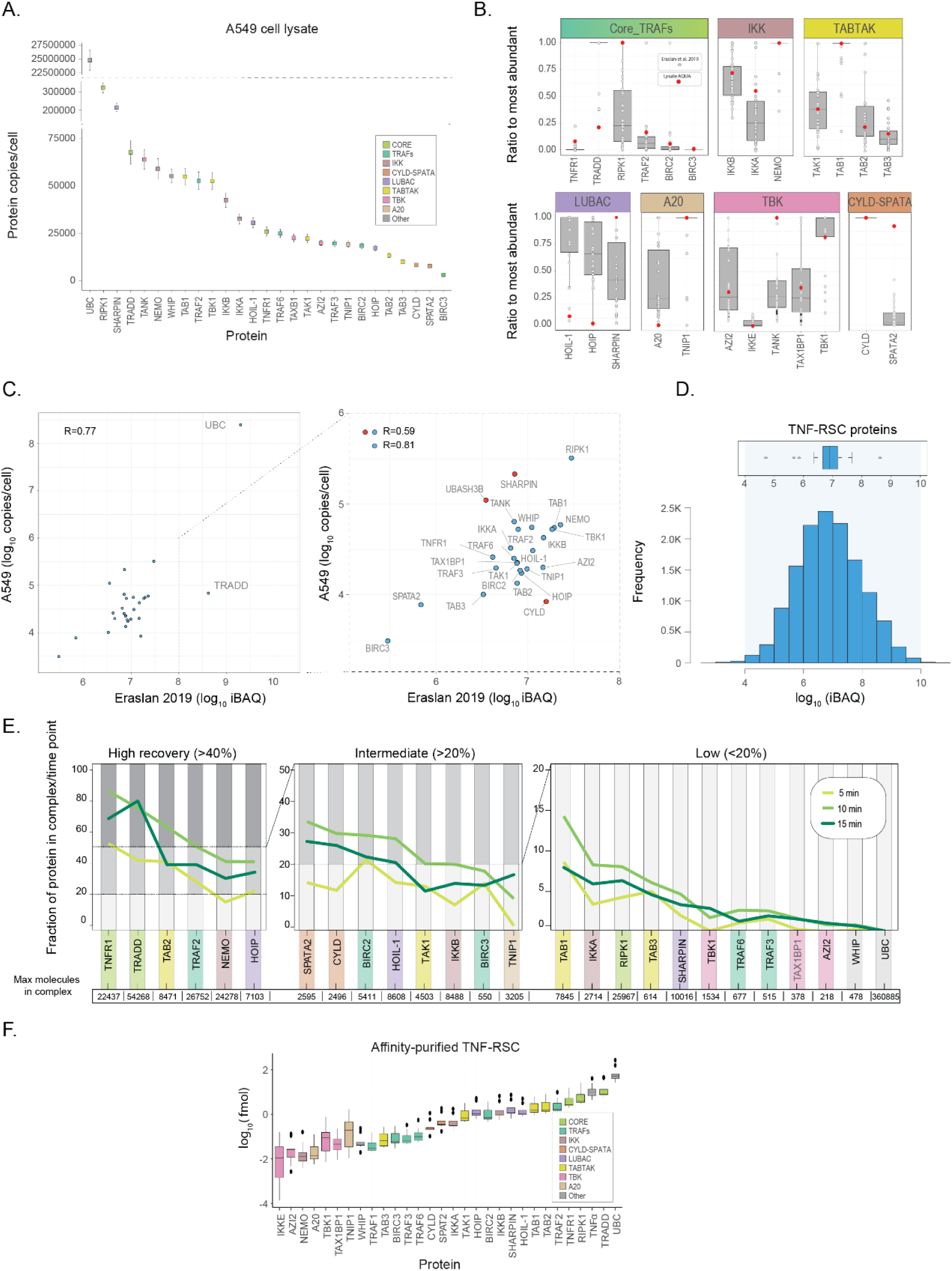
Abundance and copy/number of TNF-RSC members in lysate and affinity-purified samples. **(A)** Distribution of number of copies/cell for the indicated TNF-RSC members as estimated in the lysate-AQUA experiment. (**B)** Relative expression levels of the indicated TNF-RSC proteins as estimated in A549 (red dot) are broadly reflected in average expression levels across 29 human tissues, as determined in (*17*), normalized for the most abundant complex component. (**C)** Expression level correlation between our average estimated TNF-RSC members abundance values from A549 lysates and average values from (*17*). **(D)** Abundance distribution of TNF-RSC members based on iBAQ estimate across 29 human tissues (*17*) indicates most proteins have average expression levels. (**E)** Calculated fraction of cytosolic proteins recovered in our AP-AQUA-MS experiment across the indicated time points (yield of purification). (**F)** Distribution of amount (fmol) of TNF-RSC members in the AP-AQUA-MS experiment.

